# VenusPheno: Transfer-Guided Task Grouping for Multi-Task Bacterial Phenotype Prediction

**DOI:** 10.64898/2026.11.24.768817

**Authors:** Jinling Yuan, Bingxin Zhou, Liang Hong

## Abstract

Understanding how genomic information gives rise to bacterial phenotypes is a fundamental problem in biology. Genome-level prediction requires models that integrate information from thousands of proteins, while related phenotypes may share genomic signals. However, not all phenotype tasks benefit from being learned to-gether, and suitable task groups can be difficult to define from biological knowledge alone. We present VenusPheno, a framework that combines multiple-instance learning over pretrained ProtT5 protein embeddings with transfer-guided multi-task learning. An attention-based Set Transformer aggregates the proteins in each genome into a genome-level representation. We evaluate VenusPheno on 14 binary phenotype tasks from 12,178 bacterial genomes using genus-disjoint splits. Under single-task training, VenusPheno outperforms computational, conventional machine-learning, and deep-learning baselines in terms of mean balanced accuracy. We then use leave-one-task-out analysis to estimate directed transfer relationship between phenotypes, revealing both beneficial and detrimental interactions. Grouping tasks based on these transfer relationships further improves mean balanced accuracy over independent training and joint training of all tasks. Although inferred without biological priors, the resulting groups are consistent with biologically plausible relationships among phenotypes. VenusPheno therefore combines accurate phenotype prediction with a data-driven strategy for determining which tasks should be learned together. The PyTorch implementation is publicly available at https://github.com/YJinling/VenusPheno.

## 1 Introduction

Understanding the relationship between genotype and phenotype is a fundamental problem in biology [1]. In bacteria, measurable phenotypes emerge from the combined effects of genes and proteins [2,3]. These proteins interact and work together in multiple modules and pathways, ultimately shaping the properties of the cell as a whole, such as motility, sporulation, and carbon source utilization. Understanding how genomic information gives rise to these phenotypes helps characterize bacterial functions and explain phenotypic diversity [4].

Experimental measurements provide direct evidence of bacterial physiological capabilities and help establish links between genotypes and phenotypes. However, annotating bacterial phenotypes experimentally is labor-intensive and difficult to scale across diverse strains and traits [5]. Meanwhile, advances in sequencing technologies have generated an unprecedented volume of bacterial genomic data, yet many sequenced genomes lack corresponding experimental phenotype annotations [5,6]. This gap between genomic and phenotypic data limits our ability to identify bacteria with useful physiological traits and hinders the investigation into the genetic basis of these traits. To help bridge the gap, computational approaches have been developed to infer bacterial phenotypes from genomic information, extending phenotype annotation to strains lacking experimental measurements.

Computational approaches for bacterial phenotype prediction have evolved from knowledge-based methods to feature-based machine learning and, more recently, deep learning methods. Knowledge-based methods infer phenotypes from established genotype–phenotype relationships, and their applicability depends on the development of such knowledge [7,8]. Feature-based machine learning methods train phenotype predictors using genomic features such as protein-family annotations [5,9] and *k*-mers [10,11]. Although effective in some settings, protein-family features strongly depend on predefined functional categories and the completeness of annotations, while *k*-mer-based approaches represent genomic sequences as short fragments without explicitly preserving proteins as functional units. However, these feature-based representations, which summarize genomic information through predefined protein families or short sequence fragments, may not capture the connections between genomic components and cellular functions in the most explicit and direct manner. Since proteins are the direct molecular effectors of many cellular functions, modeling them as functional units provides a biologically grounded approach to genotype–phenotype prediction. More recently, pretrained models such as Bacformer [12] and Evo 2 [13] have demonstrated the ability to learn rich representations from large-scale genomic sequence data. These models can be adapted to downstream phenotype prediction through supervised learning. However, when pretrained genome-level representations are used as fixed features, downstream phenotype labels cannot directly influence how these representations are constructed. As a result, the relative importance of individual proteins in the genome representation cannot be adjusted according to their relevance to different phenotype tasks. This motivates a learning framework in which phenotype supervision can guide the aggregation of protein-level information into genome-level representations. Together, these considerations highlight two complementary challenges: (1) **how to integrate protein-level information for organism-level phenotype prediction**, and (2) **how to effectively leverage organism-level phenotypic supervision to guide the learning of underlying protein-level patterns**.

These cross-scale challenges show a requirement of linking microscopic elements (*e*.*g*., proteins) to macroscopic labels (*e*.*g*., phenotypes). In bacteria, different phenotypes characterize distinct aspects of the same organism and are associated with a common set of genome-encoded proteins, some of which may contribute to multiple phenotypes through shared molecular functions and pathways [4]. This shared molecular basis provides a biological rationale for multi-task learning (MTL), where different phenotype labels may offer complementary supervision for learning protein-level patterns relevant to multiple phenotypes. More broadly, this represents a class of learning problems in which each sample consists of a set of entities and is associated with multiple prediction tasks. Models may therefore benefit from both the information contained in individual entities and the supervision shared across tasks. However, whether joint learning consistently improves prediction, or whether including more tasks necessarily brings greater benefits, especially in microbial phenotype prediction, remains unclear. Previous studies in computer vision and multi-task optimization have investigated task relationships through transfer learning [14], empirical joint-training performance [15], and loss- or gradient-based analyses [16,17]. These studies suggest that task interactions can be heterogeneous, with transfer effects varying in direction and magnitude, and that seemingly related tasks may not necessarily benefit from joint training. Yet, such task-transfer relationships remain relatively unexplored in bacterial phenotype prediction, where diverse physiological traits arise from the same genome and may share underlying molecular determinants. This raises two intriguing questions: whether different bacterial phenotype tasks influence one another under joint supervision, and, if so, how these phenotypes influence one another when learned jointly. **A central methodological challenge here is to identify these heterogeneous task-transfer relationships from data and selectively exploit beneficial supervision to improve prediction across tasks**.

To address these challenges, we introduce VenusPheno, a framework that integrates bottom-up protein information through multiple-instance learning with top-down joint phenotypic supervision through multi-task learning (Fig. 1A). VenusPheno uses a shared multiple-instance learning architecture under different training configurations: single-task learning (STL), which trains each phenotype independently, and multi-task learning (MTL), which jointly optimizes a shared genome representation across multiple phenotype tasks while retaining task-specific prediction heads. To determine which tasks should be learned together, we introduce a transfer-guided grouping strategy based on leave-one-task-out (LOTO) analysis, yielding grouped MTL as our final training configuration.

**Fig. 1.**
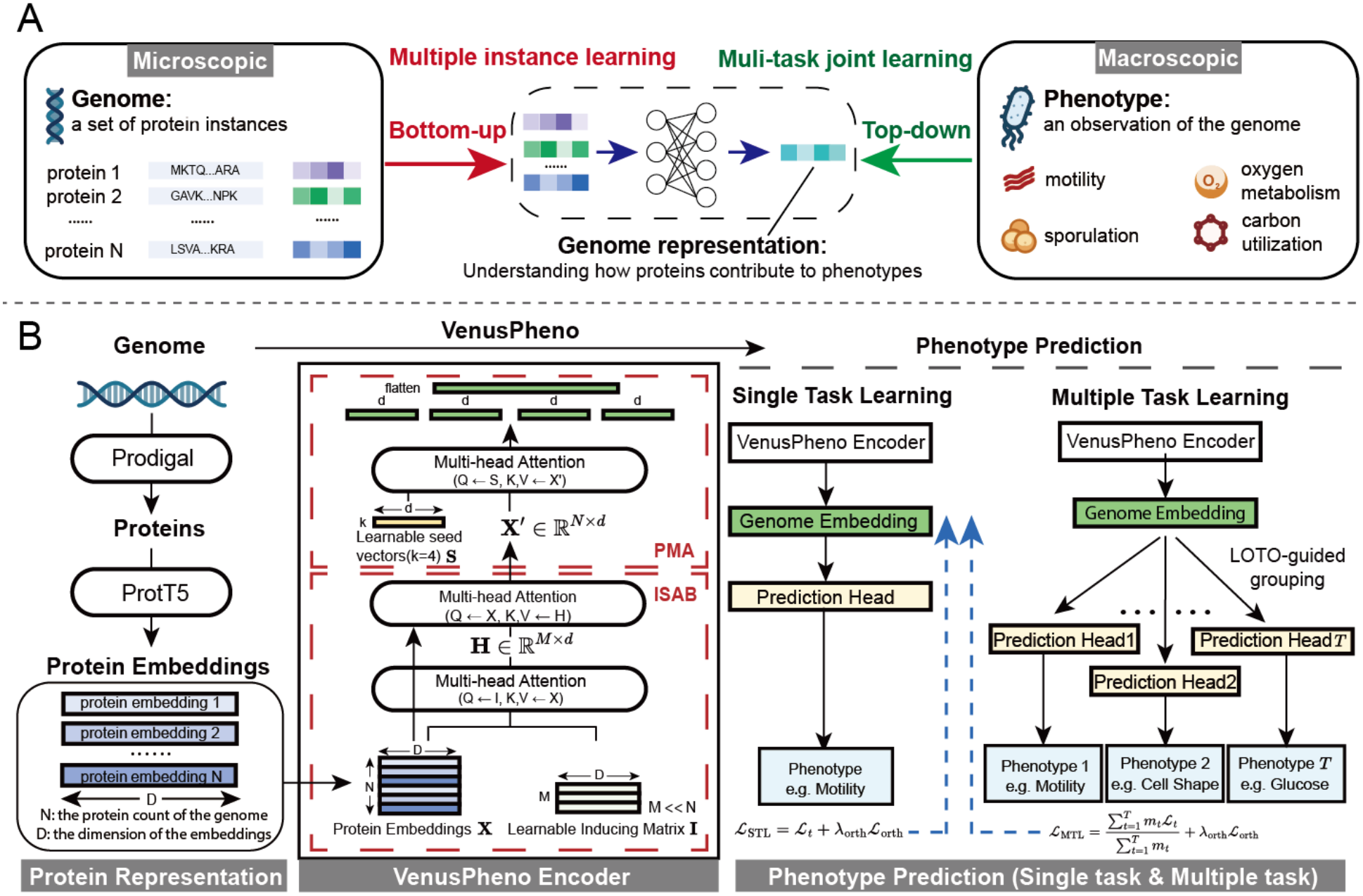
Conceptual overview and architecture of VenusPheno. (A) Bottom-up and top-down learning perspective. A bacterial genome is represented as a set of protein instances, whose information is aggregated through multiple instance learning to predict organism-level phenotypes. Different phenotypes capture complementary aspects of the organism and can provide top-down joint supervision to guide the learning of phenotype-relevant protein representations. (B) VenusPheno architecture. The architecture consists of three stages. First, in representation stage, proteins predicted by Prodigal are encoded into embeddings using ProtT5. Second, in VenusPheno encoder, a Set Transformer-based aggregation module combines Induced Set Attention Blocks (ISAB) with Pooling by Multihead Attention (PMA), using four learnable seed vectors to obtain genome-level representations. Third, in the prediction stage, the resulting representations are used for STL, MTL or LOTO-guided grouped MTL, with task-specific prediction heads.

As illustrated in Fig. 1B, each bacterial genome is represented as a variable-sized set of protein embeddings derived from a protein language model. VenusPheno then employs a set Transformer encoder [18] consisting of Induced Set Attention Blocks (ISAB) and Pooling by Multihead Attention (PMA), enabling the integration of information across proteins. Multiple PMA seeds, together with orthogonality regularization, are also incorporated to encourage complementary representations that may capture heterogeneous functional information. The resulting genome-level representations are used for phenotype prediction under either STL or MTL. We further introduce a transfer-guided task grouping strategy to determine which phenotypes should be learned jointly. To characterize task relationships and leverage the transfer structure, we perform LOTO analysis [19] to quantify how excluding one task from joint training affects the performance of others, constructing a directed task-transfer graph. We then use this graph to guide task grouping and improve the multi-task phenotype prediction. Finally, we analyze the PMA attention of STL and MTL to identify biologically meaningful functional patterns and explore potential shared signals captured by joint supervision.

Our study makes three main contributions. First, we demonstrate that integrating genome-wide protein information through multiple-instance learning provides an effective approach to organism-level phenotype prediction. VenusPheno achieves strong predictive performance across diverse bacterial phenotypes, with further improvements obtained through multi-task supervision. Second, we reveal heterogeneous and directional task-transfer relationships among bacterial phenotypes, including both beneficial and detrimental interactions. By identifying these relationships through LOTO analysis and using them to guide task grouping, we further improve predictive performance over both single-task learning and full-task MTL, demonstrating the value of selective joint supervision. Third, we show that VenusPheno captures biologically relevant protein-level signals associated with bacterial phenotypes. Attention analysis identifies phenotype-associated protein families consistent with known cellular functions, while comparisons between STL and grouped MTL reveal changes in these attention patterns, providing insights into how joint supervision shapes the protein-level information emphasized by the model.

## 2 Methods

### 2.1 Dataset and experimental setup

We assembled a genome-phenotype dataset using phenotype annotations and their corresponding genome assembly accessions from the publicly available Bacformer dataset [12]. Bacterial genome assemblies were retrieved from NCBI RefSeq and GenBank using these accessions [20,21]. Following taxonomic filtering, the final dataset comprised 12,178 bacterial genomes spanning 2,061 genera, with phenotype annotations covering 14 binary classification tasks. These tasks included motility, sporulation, Gram staining, two cell shape categories (bacillus and coccus), three oxygen metabolism categories (aerobic, anaerobic, and facultative), and six carbon utilization traits (citrate, glucose, glycerol, lactose, maltose, and sucrose). As not all genomes had annotations for every phenotype, the number of labeled samples varied across tasks. Detailed task definitions and sample distributions are provided in Supplementary Table S1.

To assess generalization to previously unseen bacterial genera, we adopted a genus-disjoint data split. Closely related bacteria often share genomic and phenotypic characteristics, potentially introducing phylogenetic confounding in phenotype prediction. We therefore assigned entire genera, rather than individual genomes, to the training, validation, or test set, ensuring that no genus represented in the test set appeared in the training set. We selected a genus-disjoint partition to approximate the target 7:1:2 ratio while balancing genome counts and task-specific class distributions. The resulting training, validation, and test sets contained 8,409, 1,257, and 2,512 genomes, respectively, spanning 1,443, 206, and 412 mutually exclusive genera. Detailed partition optimization and validation procedures are provided in Supplementary Section A.2.

We compared VenusPheno against machine learning and deep learning baselines, including Traitar [5], PhenoTrex-XGB [9], BacML (RF, HGBM, RNN, and CNN) [10], BacteriaNet [22], Bacformer [12], and Evo 2 [13]. These methods represent genomes using diverse features, including protein functional annotations, *k*-mer frequencies, ge-nomic sequences, and pretrained sequence embeddings. We used balanced accuracy (BAcc), defined as the average of sensitivity and specificity, as the primary evaluation metric to account for class imbalance. Detailed data processing, splitting procedures, baseline implementations, and evaluation settings are provided in Supplementary Section A.

### 2.2 Model architecture

As illustrated in Fig. 1B, VenusPheno consists of three sections: protein representation, VenusPheno encoder, and phenotype prediction.

#### Protein Representation

In the first stage, protein-coding genes were predicted from each bacterial genome using Prodigal (v2.6.3) [23]. The resulting protein sequences were encoded using the pretrained protein language model ProtT5-XL-UniRef50 [24]. Residue-level embeddings were mean-pooled to obtain a 1024-dimensional representation for each protein. Each genome was thus represented as a variable-sized set of protein embeddings, 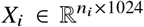, where *n*_*i*_ denotes the number of proteins in genome *i*.

#### VenusPheno Encoder

In the second stage, VenusPheno aggregates protein-level embeddings into a genome-level representation using a multiple-instance learning framework. We use an attention-based set encoder inspired by the Set Transformer architecture [18]. For a genome represented by a protein embedding matrix **X** ∈ ℝ^*N*×*D*^, where *N* is the number of proteins in the genome and *D* = 1024 is the embedding dimension, the encoder applies an Induced Set Attention Block (ISAB) followed by Pooling by Multihead Attention (PMA).

ISAB models contextual relationships among proteins through a set of *M* = 64 learnable inducing points **I** ∈ ℝ^*M*×*D*^. Rather than computing pairwise attention across all *N* proteins, ISAB performs two successive attention operations:

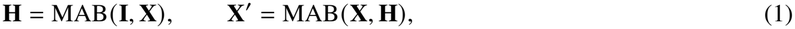

where MAB denotes a multihead attention block, with its first argument serving as the query and its second as the keys and values. The inducing representations **H** ∈ ℝ^*M*×*D*^ first aggregate information from the protein set, which is subsequently propagated back to individual proteins to obtain context-aware representations **X**^′^∈ ℝ^*N*×*D*^. This design reduces the attention complexity from *O N*^2^ to *O N M* , enabling efficient modeling of genome-wide protein contexts.

PMA then aggregates these contextualized protein representations into a fixed-size genome representation using *K* = 4 learnable seed vectors **S** ∈ ℝ^*K*×*D*^:

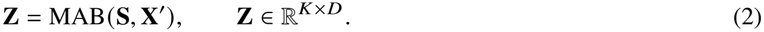

Each seed independently attends to the protein set, allowing the encoder to extract multiple potentially distinct patterns of protein-level information. To encourage complementary representations and reduce redundancy across seeds, we introduce an orthogonality regularization term that penalizes pairwise cosine similarities between the PMA outputs:

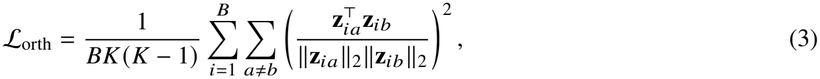

where *B* is the batch size and **z**_*ia*_ denotes the representation produced by seed *a* for genome *i*. The four resulting representations are concatenated into a fixed-length genome embedding for downstream phenotype prediction. Padding masks are applied during protein-level attention to accommodate variable-sized protein sets.

#### Phenotype Prediction

In the third stage, the concatenated genome representation is passed through a residual multilayer perceptron, followed by a task-specific linear classification head. VenusPheno supports both STL and MTL using the same architecture. In STL, each phenotype task is trained independently with its own encoder and prediction module. In MTL, tasks share the encoder and residual MLP while maintaining separate prediction heads, allowing multiple phenotype labels to jointly guide representation learning.

Both settings use class-weighted binary cross-entropy to account for class imbalance, together with the orthogonality regularization defined above. Their training objectives are formulated as:

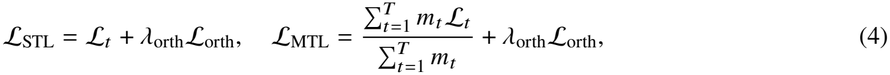

where *ℒ*_*t*_ denotes the class-weighted BCE loss for task *t, T* is the number of jointly trained tasks, and *m*_*t*_ indicates whether valid labels are available for task *t* in the current batch. The regularization coefficient *λ*_orth_ is set to 0.01. Detailed training configurations are provided in Supplementary Table S2.

### 2.3 LOTO-guided task grouping

To quantify task interactions under joint supervision, we employed a leave-one-task-out (LOTO) strategy. We first trained a full-task MTL reference model using all 14 phenotype tasks. Then we independently retrained the same MIL architecture 14 times, each time excluding one task while retaining the remaining 13 tasks. The directional transfer effect from task *i* to task *j* was defined as:

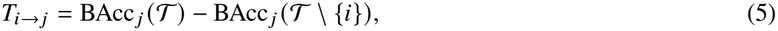

where *T* denotes the complete task set and BAcc_*j*_ represents the validation balanced accuracy for task *j*. Positive and negative values indicate beneficial and detrimental transfer, respectively. The resulting directed transfer matrix captures potentially asymmetric interactions between tasks.

We then used the LOTO transfer effects to guide task partitioning, aiming to preserve beneficial transfer while limiting detrimental interactions. For a partition *G*, the net incoming transfer to each task *j* was defined as *u* _*j*_ (*G*) =Σ_*i*∈*G*(*j*), *i*≠ *j*_ *T*_*i*→*j*_ , where *G*(*j*) denotes the group containing task *j*. We further defined the total net transfer as

*U*(G) = Σ_*j*_ *u* _*j*_ (*G*) and the negative-transfer burden as *H*(*G*) = Σ_*j*_ max(™*u* _*j*_ (*G*), 0). The grouping objective was formulated as:

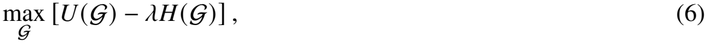

where *λ* controls the penalty on negative transfer. We exhaustively searched feasible partitions, assigning each task to exactly one group and constraining group sizes to three to six tasks. We examined *λ* ∈ {0.5, 1, 2} to assess the sensitivity of task assignments to the negative-transfer penalty. The resulting group assignments were consistent across the examined settings for each group count. To characterize the optimization landscape, we further evaluated candidate partitions in terms of their total net transfer and negative-transfer burden, and identified Pareto-optimal solutions representing trade-offs between these two quantities (Fig. 2A). We selected the optimal three-group partition for subsequent grouped MTL training.

**Fig. 2.**
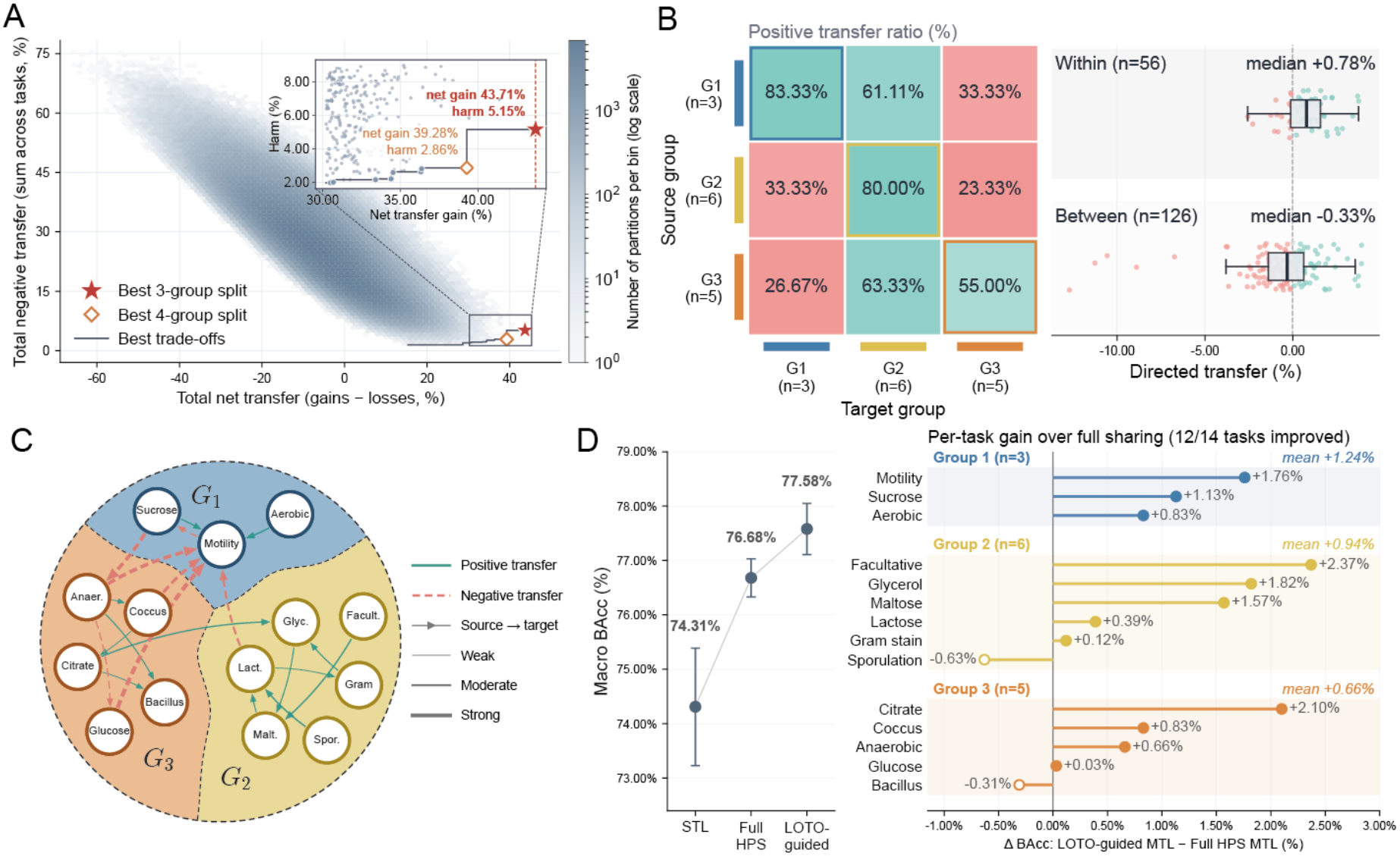
LOTO-guided task grouping captures directional transfer relationships and improves multi-task prediction. (A) Optimization landscape of candidate partitions with 3-6 tasks per group, characterized by total net transfer and target-level negative-transfer burden. Pareto-optimal solutions and the selected three-group (red star) and four-group (orange diamond) partitions are highlighted, with an inset showing the selected solutions. (B) Directed transfer patterns among the three selected groups. The heatmap shows the percentage of positive transfer edges between source-target groups. The right panel summarizes within- and between-group transfer effects by medians and interquartile ranges. (C) Directed transfer network of the selected three-group partition. Shaded regions indicate task groups; solid teal and dashed red arrows represent positive and negative transfer. Arrow direction indicates source-to-target transfer, and width scales with absolute transfer magnitude (|*T*|). For visualization, each group retains one edge per task, selected to preserve connectivity and prioritize high-magnitude interactions, while the six strongest cross-group edges are shown. (D) Performance comparison of VenusPheno-STL, VenusPheno-MTL, and VenusPheno-LOTO. The left panel shows BAcc, and the right shows per-task changes relative to VenusPheno-MTL, grouped by the partition. VenusPheno-LOTO improves macro BAcc from 76.68% to 77.58%, with gains in 12 out of 14 tasks. Differences are in percentage points (displayed using % notation).

Following task partitioning, an independent VenusPheno MTL model was trained for each group, with shared representations learned among tasks within the same group. Predictions from all groups were combined for evaluation across the 14 phenotype tasks.

### 2.4 Interpretability analysis

To investigate the protein-level signals learned by VenusPheno, we developed an attention-based functional analysis workflow (Fig. 3A). Interpretability analyses were performed on the held-out validation set rather than on the full collection of bacterial genomes. For each phenotype, we retained genomes with an observed label for that phenotype and a successfully exported PMA attention profile. No genomes were selected according to their functional annotations or attention values before this eligibility filtering. For matched comparisons between STL and MTL models, we used the intersection of eligible validation genomes, such that both models were evaluated on exactly the same genomes for each phenotype.

**Fig. 3.**
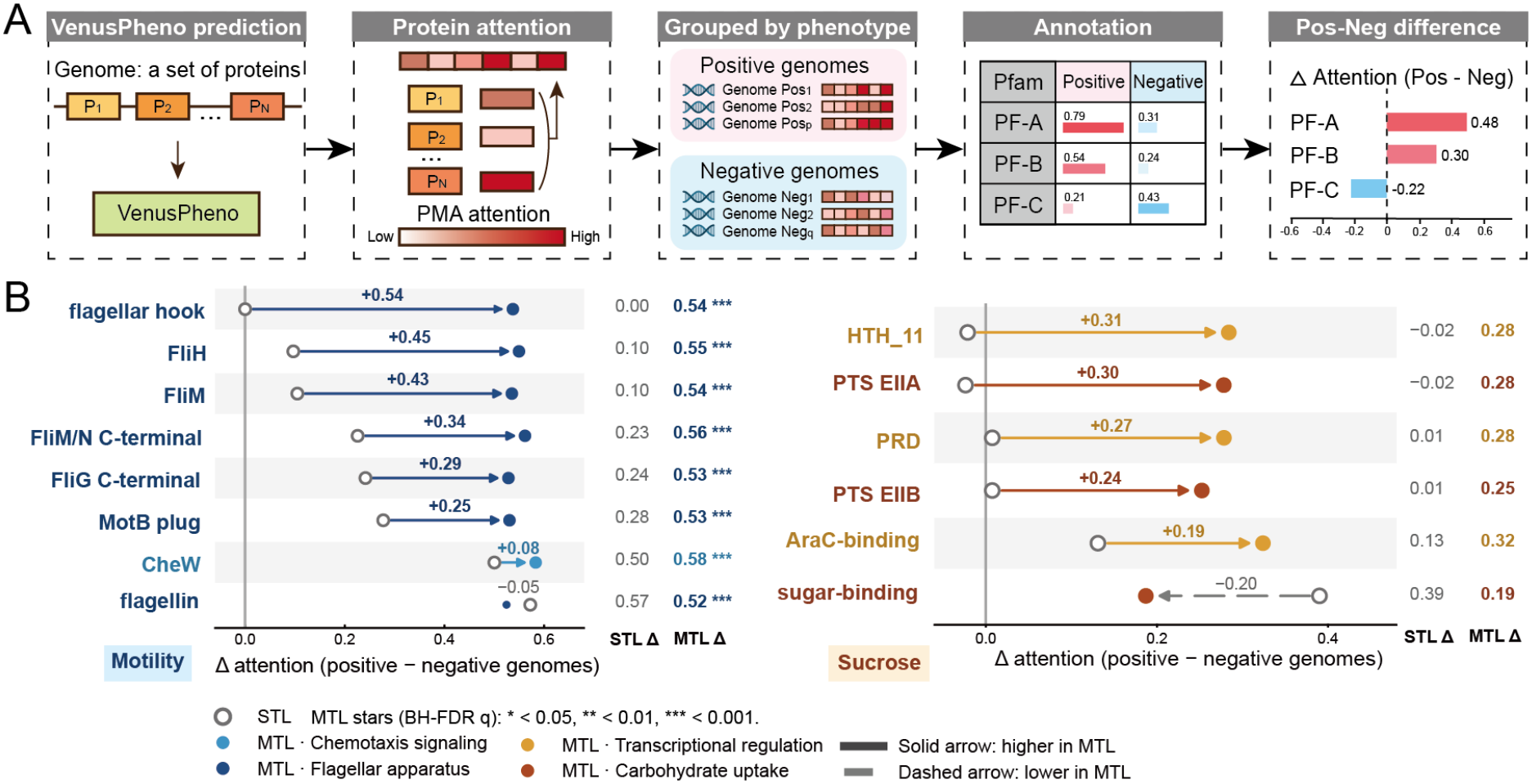
Phenotype-associated protein attention patterns under single-task and multi-task learning. (A) Workflow for protein-level attention analysis. Protein attention scores from VenusPheno are aggregated by Pfam family and compared between phenotype-positive and phenotype-negative genomes. The attention contrast (Δ attention) is calculated as the difference in mean family-level attention between the two groups. (B) Phenotype-associated attention differences in phenotype-related protein families under STL and MTL, shown for motility (left) and sucrose utilization (right). For motility, MTL yields larger attention differences across proteins involved in flagellar assembly, motor function, and chemotaxis. For sucrose utilization, increased differences are observed in protein families associated with phosphotransferase system transport and transcriptional regulation. Open circles represent STL, filled circles represent MTL, and arrows indicate the direction of change from STL to MTL. Colors distinguish functional categories. Numbers above arrows indicate Δ_MTL_ ™ Δ_STL_, while the right columns show the corresponding attention differences for each model. Stars indicate significant positive–negative differences under MTL after BH-FDR correction (*q* < 0.05, *q* < 0.01, *q* < 0.001).

We extracted protein-level attention weights from the PMA layer of the trained models. Each genome yielded attention distributions across all PMA head–seed slots (four pooling seeds and four attention heads). Attention weights were ranked independently within each slot to account for differences in protein-bag size and the softmax normalization of PMA weights. For the functional attention analysis, the top 5% of proteins within each slot were designated as highly attended instances. These proteins were mapped to functional families using eggNOG annotations [25,26]. For each genome and functional family, we calculated a functional attention score as the fraction of PMA slots in which that family appeared among the highly attended proteins.

To identify phenotype-associated functional patterns, we compared functional attention scores between phenotype-positive and phenotype-negative genomes in the validation set. The attention contrast (Δ) for each family was defined as the difference in mean functional attention scores between the two groups. We performed the same analysis using matched STL and MTL models on identical genomes, and quantified the effect of joint supervision as Δ_MTL_ ™ Δ_STL_. Statistical significance was assessed using two-sided Welch’s tests with Benjamini–Hochberg false discovery rate correction.

For visualization of recurrent high-attention proteins and pathway enrichment, we analyzed all eligible phenotype-positive validation genomes for each focal task. Proteins were ranked by their mean rank across PMA slots, and the top 10% within each genome were retained for this descriptive analysis. Recurrently selected proteins were summarized by eggNOG gene name. KEGG pathway enrichment was tested against all other annotated proteins from the same set of phenotype-positive genomes using one-sided Fisher’s exact tests, followed by Benjamini–Hochberg correction. Thus, the pathway panels summarize model attention within the relevant positive validation cohort and do not represent a genome-wide or test-set-selected analysis.

## 3 Results

### 3.1 VenusPheno achieves competitive performance across bacterial phenotypes, with heterogeneous gains from multi-task learning

To evaluate the effectiveness of our model design, we conducted a comparative benchmark across 14 binary classification tasks. These tasks cover diverse aspects of bacterial biology, including cell structure, motility and sporulation, oxygen metabolism, and carbon utilization, representing biological processes with potentially distinct genomic determinants. We developed three training configurations of VenusPheno: VenusPheno-STL (single-task learning, where each phenotype is trained independently), VenusPheno-MTL (full-task multi-task learning, where all 14 phenotypes jointly optimize a shared model backbone), and VenusPheno-LOTO (LOTO-guided grouped multi-task learning, where phenotype tasks are grouped according to their estimated directional transfer effects). We compared these configurations against machine-learning and deep-learning baselines using diverse genome representations, including protein annotation-derived features [5,9], *k*-mer representations [10], genomic sequences [22], and pretrained sequence representations [12,13]. All experiments followed a genus-disjoint evaluation protocol to assess generalization to un-seen bacterial genera and reduce potential bias arising from taxonomic similarity between training and test genomes (Section 2.1). The results are presented in Table 1.

**Table 1.** Performance comparison of bacterial phenotype prediction methods across 14 tasks. Balanced accuracy (BAcc, %) is reported as mean ± std over 3 independent runs where available. N/A marks phenotypes not supported by the method.

| **Model** | **Macro<br>BAcc (%)** | **Morphology** |  |  |  |  | **Metabolism** |  |  | **Carbon utilization** |  |  |  |  |  |
|---|---|---|---|---|---|---|---|---|---|---|---|---|---|---|---|
|  |  | Motility | Sporulation | Gram | Cell shape<br>(Bacillus) | Cell shape<br>(Coccus) | Aerobic | Anaerobic | Facultative | Citrate | Glucose | Glycerol | Lactose | Maltose | Sucrose |
| **Machine learning methods** |  |  |  |  |  |  |  |  |  |  |  |  |  |  |  |
| *Protein annotation-derived features* |  |  |  |  |  |  |  |  |  |  |  |  |  |  |  |
| Traitar | 73.89 | 80.46 | 89.78 | 97.32 | 81.66 | 84.56 | 73.96 | 91.47 | 83.57 | 54.48 | 52.77 | 53.52 | 61.54 | 62.21 | 67.17 |
| PhenoTrex-XGB | 67.79<br>±0.08 | **86.61**<br>±0.10 | 95.41<br>±0.36 | **97.15**<br>±0.05 | 54.75<br>±0.49 | 54.60<br>±0.88 | 65.80<br>±1.02 | 78.08<br>±0.37 | 68.92<br>±0.67 | 51.22<br>±0.58 | 60.44<br>±0.51 | 50.11<br>±0.29 | 56.84<br>±0.42 | **62.67**<br>±1.17 | **66.44**<br>±0.56 |
| *k-mer-derived features* |  |  |  |  |  |  |  |  |  |  |  |  |  |  |  |
| BacML-RF | 59.22<br>±0.25 | 75.08<br>±1.54 | 73.45<br>±0.46 | 86.56<br>±2.33 | 53.53<br>±0.17 | 50.50<br>±0.20 | 57.91<br>±0.79 | 68.47<br>±1.02 | 53.64<br>±0.74 | 49.89<br>±0.00 | 53.44<br>±1.05 | 50.61<br>±0.20 | 50.25<br>±0.09 | 53.07<br>±0.48 | 52.74<br>±1.26 |
| BacML-HGBM | 64.55<br>±0.17 | 76.01<br>±0.19 | 84.85<br>±0.24 | 88.57<br>±0.59 | 55.62<br>±0.14 | 51.28<br>±0.14 | 65.81<br>±0.31 | 78.29<br>±0.25 | 69.77<br>±0.19 | 58.22<br>±0.55 | 55.06<br>±0.76 | 50.58<br>±2.27 | 55.46<br>±0.62 | 56.74<br>±1.23 | 57.43<br>±1.41 |
| **Deep learning methods** |  |  |  |  |  |  |  |  |  |  |  |  |  |  |  |
| *Nucleotide sequence* |  |  |  |  |  |  |  |  |  |  |  |  |  |  |  |
| BacteriaNet | 59.22 | 63.90 | 69.23 | 74.14 | 50.72 | 69.89 | 50.00 | 47.03 | 48.88 | N/A | N/A | N/A | N/A | N/A | N/A |
| *k-mer-derived features* |  |  |  |  |  |  |  |  |  |  |  |  |  |  |  |
| BacML-RNN | 65.83<br>±1.42 | 70.09<br>±7.34 | 89.67<br>±1.99 | 90.00<br>±1.90 | 52.57<br>±3.53 | 70.13<br>±13.39 | 65.66<br>±0.73 | 82.14<br>±0.93 | 80.57<br>±0.42 | 60.03<br>±1.41 | 52.63<br>±2.04 | 48.68<br>±0.72 | 56.48<br>±1.55 | 53.02<br>±0.60 | 50.00<br>±0.00 |
| BacML-CNN | 65.46<br>±1.56 | 72.86<br>±1.81 | 83.54<br>±3.86 | 87.13<br>±2.74 | 57.57<br>±8.76 | 61.48<br>±13.84 | 64.03<br>±0.86 | 75.11<br>±2.20 | 79.70<br>±0.39 | 63.92<br>±0.25 | 51.61<br>±1.13 | 50.14<br>±0.39 | 58.39<br>±1.24 | 53.64<br>±1.01 | 57.32<br>±0.45 |
| *Pretrained sequence representations* |  |  |  |  |  |  |  |  |  |  |  |  |  |  |  |
| Bacformer | 73.77<br>±0.54 | 85.11<br>±0.19 | 94.99<br>±0.17 | 96.25<br>±0.04 | 65.67<br>±6.93 | 79.93<br>±0.27 | 69.79<br>±0.76 | 92.23<br>±0.21 | **84.77**<br>±0.06 | 66.18<br>±0.49 | 55.20<br>±2.00 | 55.67<br>±0.85 | 62.84<br>±1.18 | 59.23<br>±1.09 | 64.94<br>±0.58 |
| Evo2 | 64.61<br>±0.84 | 56.98<br>±5.39 | 90.06<br>±2.05 | 81.53<br>±2.25 | 77.11<br>±0.26 | 78.28<br>±1.06 | 65.43<br>±0.70 | 77.45<br>±5.91 | 67.99<br>±6.89 | 52.01<br>±2.85 | 51.89<br>±1.24 | 49.73<br>±0.44 | 54.93<br>±0.85 | 50.49<br>±2.18 | 50.71<br>±1.01 |
| **Our model** |  |  |  |  |  |  |  |  |  |  |  |  |  |  |  |
| VenusPheno-STL | 74.31<br>±1.08 | 86.93<br>±0.54 | 96.40<br>±0.20 | 96.88<br>±0.28 | 68.34<br>±13.15 | 88.72<br>±1.73 | 70.03<br>±0.40 | 91.46<br>±1.57 | 81.30<br>±2.51 | 63.08<br>±5.01 | 58.50<br>±2.44 | 56.36<br>±0.72 | 61.04<br>±2.11 | 58.15<br>±1.23 | 63.12<br>±2.29 |
| VenusPheno-MTL | **76.68**<br>±0.35 | 84.63<br>±0.14 | **95.59**<br>±0.35 | 96.98<br>±0.13 | 84.87<br>±2.91 | **89.90**<br>±0.61 | **75.56**<br>±0.50 | **93.00**<br>±0.55 | 82.58<br>±1.47 | 63.70<br>±2.86 | 58.61<br>±0.93 | **57.05**<br>±0.37 | **64.27**<br>±1.45 | 61.66<br>±0.71 | 65.07<br>±0.98 |
| VenusPheno-LOTO | 77.58<br>±0.47 | 86.39<br>±0.90 | 94.96<br>±2.66 | 97.10<br>±0.39 | **84.56**<br>±0.97 | 90.73<br>±1.10 | 76.39<br>±1.42 | 93.66<br>±0.55 | 84.95<br>±0.07 | **65.80**<br>±2.08 | **58.64**<br>±0.91 | 58.87<br>±0.88 | 64.66<br>±0.95 | 63.23<br>±3.15 | 66.20<br>±1.10 |

† The top three are highlighted by First, **Second**, Third.

Overall, VenusPheno achieved strong predictive performance across diverse bacterial phenotypes, with multi-task supervision further improving aggregate performance despite heterogeneous gains across tasks.

VenusPheno achieved the highest overall macro-averaged BAcc among the evaluated methods while maintaining broad predictive competitiveness across diverse bacterial phenotypes (Table 1). Under single-task learning, VenusPheno-STL outperformed the strongest external baselines, Traitar and Bacformer, demonstrating the effectiveness of protein-level multiple instance representations for organism-level phenotype prediction. Beyond aggregate performance, considering all three VenusPheno configurations, at least one configuration achieved the highest mean BAcc on 10 of the 14 tasks and ranked among the top three methods for every evaluated phenotype. These competitive results spanned cellular traits, oxygen metabolism, and carbon utilization, indicating consistent performance across distinct biological processes. In comparison, external baseline methods exhibited more pronounced task-dependent strengths and weaknesses. For example, protein annotation-derived approaches performed well on Gram staining but were less competitive on citrate utilization, whereas Bacformer performed strongly on certain metabolic traits but showed weaker performance on cell shape prediction. Together, these findings demonstrate the broad predictive capability of VenusPheno across bacterial phenotypes with diverse genomic determinants.

Importantly, we found that joint supervision improved prediction performance across most tasks, but its effects varied across phenotype tasks. Compared with VenusPheno-STL, VenusPheno-MTL increased macro-averaged BAcc and improved prediction on 12 of the 14 tasks (Table 1). Particularly large improvements were observed for bacillus cell shape (+16.53%) and aerobic growth (+5.53%), whereas performance declined for motility and sporulation. These contrasting outcomes reveal that the effects of multi-task supervision vary in both direction and magnitude across phenotype tasks. As phenotype tasks jointly optimize a shared backbone, such heterogeneity may arise from beneficial or detrimental interactions between tasks during training. This raises the question of how individual tasks influence one another during joint training and whether these transfer relationships can guide task grouping to improve multi-task learning.

### 3.2 Bacterial phenotype tasks exhibit structured transfer relationships that help improve multi-task learning

Though full-task MTL improved overall predictive performance, the gains varied across phenotype tasks, raising the question of how individual tasks influence one another during joint training. To characterize these relationships, we performed LOTO analysis, measuring how excluding each task from joint training affected the predictive performance of the remaining tasks. Among the 182 directed task pairs, 92 exhibited positive transfer and 89 exhibited negative transfer, and one showed no measurable transfer (Supplementary Fig. S1). The magnitude and direction of transfer varied considerably across tasks, with some tasks providing net benefits to other predictions while others introduced detrimental effects, indicating that individual tasks could either benefit or impair the prediction of other tasks under MTL. Moreover, these interactions were asymmetric, such that the effect of task *i* on task *j* could differ from that of task *j* on task *i*. Together, these findings reveal heterogeneous and directional transfer relationships among bacterial phenotype tasks, motivating a transfer-guided approach to task grouping.

To explore and leverage these interactions, we sought to identify task groupings that preserve beneficial transfer while minimizing detrimental interactions. We used the directed LOTO transfer effects to guide task partitioning, constraining each group to contain three to six tasks and optimizing the trade-off between positive and negative transfer effects. Candidate partitions were evaluated using an objective that rewards beneficial within-group transfer while penalizing detrimental transfer, with *λ* controlling the penalty strength. At *λ* = 1, the three-group partition achieved the highest estimated net transfer gain while maintaining a relatively low negative-transfer burden (Fig. 2A). We therefore selected the three-group configuration for subsequent evaluation, with the resulting task assignments illustrated in

Fig. 2C. Notably, sensitivity analysis across *λ* = 0.5, 1, and 2 yielded identical task assignments for each fixed number of groups, indicating that the identified partitions were stable across the tested penalty settings (Supplementary Fig. S2).

The selected grouping concentrated favorable transfer relationships within groups while separating many detrimental interactions. Under the partition, 71.4% of within-group directed interactions were positive, compared with 41.3% of between-group interactions. Consistently, the median transfer effect was positive within groups but negative between groups (Fig. 2B). The directed transfer network graph further illustrates this structure, with positive interactions retained within group region, while several strong negative interactions occurred across group boundaries (Fig. 2C). Notably, the resulting groups combined phenotypes from different biological categories, including morphology, oxygen metabolism, and carbon utilization, suggesting that transfer-based task relatedness extends beyond conventional phenotype classifications. Together, these findings suggest that LOTO-guided grouping can concentrate beneficial interactions within groups while reducing exposure to negative transfer.

Importantly, we found that LOTO-guided task grouping could improve overall phenotype prediction compared with full-task multi-task learning. Under genus-disjoint evaluation, the three-group configuration increased macro BAcc from 76.68% with full-task MTL to 77.58%, compared with 74.31% under single-task learning (Fig. 2D). Improvements over full-task MTL were observed in 12 of the 14 tasks, spanning all three groups. Particularly notable gains were achieved for facultative growth, citrate utilization, and motility. Although sporulation and bacillus cell shape prediction showed modest decreases, the overall improvements suggest that transfer-guided grouping can mitigate unfavorable task interactions while retaining the benefits of joint supervision.

### 3.3 PMA attention reveals functionally connected biological processes under joint phenotypic supervision

The performance improvements achieved through transfer-guided task grouping prompted us to investigate whether joint supervision enables VenusPheno to capture stronger phenotype-relevant protein signals. We extracted protein-level attention scores from the PMA module and identified highly attended proteins within each genome. These proteins were functionally characterized using eggNOG-mapper [25,26], including Pfam family assignments [27]. We first examined the occurrence of highly attended proteins across phenotype-positive genomes and performed KEGG pathway enrichment analysis [28] to assess whether the learned attention patterns reflected biologically relevant functions under STL. We then investigated how these patterns changed under joint supervision by quantifying the difference in mean attention between phenotype-positive and phenotype-negative genomes for each Pfam family (Δ attention). Statistical significance of the attention differences was assessed with BH-FDR correction (Fig. 3A). We examined motility and sucrose utilization as two representative case studies, selected based on the availability of sufficient phenotype-labeled genomes and functional annotations. Both tasks were jointly supervised alongside aerobic metabolism, providing a shared setting for investigating how joint supervision alters attention to phenotype-associated protein families.

Overall, the analysis revealed that VenusPheno identifies protein signals consistent with known phenotype-associated biological functions, while joint supervision may further highlight functionally connected biological processes across phenotypes.

Under STL, VenusPheno assigned high attention to proteins associated with known biological processes relevant to the target phenotypes. For motility, FlgK, a component of the flagellar assembly machinery, and DnaK, a molecular chaperone implicated in flagellar assembly [29], were recurrently identified among high-attention proteins. Consistently, KEGG enrichment analysis of high-attention proteins revealed strong enrichment of bacterial chemotaxis and flagellar assembly [30,31]. For sucrose utilization, recurrent high-attention proteins included carbohydrate-related proteins such as MalL [32], while KEGG analysis revealed enrichment of starch and sucrose metabolism and galactose metabolism (Supplementary Fig. S3). These findings suggest that the multiple-instance learning framework of VenusPheno can associate organism-level phenotype predictions with proteins involved in relevant cellular functions, providing protein-level biological interpretability alongside predictive performance. This further establishes a biological basis for examining how these representations change under MTL.

Under joint supervision, several functionally relevant protein families exhibited stronger phenotype-associated attention contrasts, defined as the difference in mean attention between phenotype-positive and phenotype-negative genomes (Δ attention), potentially revealing broader biological processes connecting the jointly learned phenotypes. Compared with STL, for motility, VenusPheno-LOTO increased these contrasts for flagellar hook proteins, FliH, and FliM, which are involved in flagellar assembly, protein export, and motor switching. All three showed significantly higher attention in phenotype-positive genomes under MTL after multiple testing correction. For sucrose utilization, larger positive contrasts were observed for PTS EIIA and EIIB domains, involved in carbohydrate transport and phosphorylation, and PRD and HTH_11 domains associated with transcriptional regulation. Although these differences did not reach statistical significance after multiple testing correction, they suggest changes in attention toward carbohydrate transport and regulatory functions (Fig. 3B; Supplementary Fig. S4). Interestingly, previous studies have demonstrated biological connections between carbohydrate uptake and bacterial motility, including PTS-mediated signaling to the chemotaxis machinery and sucrose-induced flagellar motility in *Bacillus subtilis* [33,34]. These findings indicate that carbohydrate utilization and motility can be functionally connected through nutrient-responsive signaling and regulatory processes. In this context, the attention changes observed under VenusPheno-LOTO suggest that joint supervision may help VenusPheno capture complementary protein-level signals involved in these biologically linked processes. More broadly, such functional connections may contribute to interactions among phenotype tasks during multitask learning, even when the tasks represent distinct organism-level traits. Protein-level attention analysis thus provides a means to investigate the biological relationships underlying task interactions beyond predictive performance alone.

## 4 Discussion and Conclusion

Predicting bacterial phenotypes from genomic sequences requires models to address two related challenges: integrating information from thousands of proteins into an organism-level representation and exploiting information shared across phenotypes without introducing detrimental task interference. VenusPheno addresses these challenges by combining a multiple-instance set encoder with transfer-guided multi-task learning. Under genus-disjoint evaluation across 14 phenotype tasks, VenusPheno achieved the highest macro-averaged balanced accuracy among the evaluated methods. Moreover, grouping tasks according to the LOTO transfer relationships improved performance over both independent single-task training and joint training of all phenotypes. These results show that phenotype supervision can provide useful information not only for individual prediction tasks, but also for learning how genome-level representations should be shared across tasks.

Our results demonstrate that phenotype labels can provide complementary supervision, but that their value depends on the structure of cross-task transfer. Full-task MTL improved overall prediction while producing uneven effects across phenotypes, and LOTO analysis revealed directed and asymmetric relationships containing both beneficial and detrimental transfer. Organizing tasks according to these relationships yielded the highest overall predictive performance, with VenusPheno-LOTO outperforming VenusPheno-STL and VenusPheno-MTL on most phenotype tasks. The resulting groups crossed conventional phenotype categories, showing that predictive compatibility can capture relationships that are difficult to define from biological knowledge alone. Protein-level attention analysis further provided a functional interpretation of these interactions: VenusPheno recovered established phenotype-associated signals under STL, while grouped supervision strengthened attention to functions related to flagellar machinery, carbohydrate transport, and transcriptional regulation. Together, these findings show that data-driven task organization can improve phenotype prediction by selectively reinforcing biologically informative signals shared across tasks.

Several limitations remain. First, phenotype annotations in existing databases are unevenly distributed, and experimentally observed phenotypes may depend on environmental conditions that are not represented by genomic sequences alone. This may partly account for the lower predictive performance on some carbon utilization tasks. Second, protein-level attention reflects the importance assigned by the model and does not establish a causal relationship between individual proteins and phenotypes. Third, the transfer relationships estimated by LOTO are specific to the current dataset, model architecture, and training setting, and their generalizability should be evaluated in other settings. Future work could incorporate environmental context, develop more generalizable task-grouping strategies, and experimentally validate the phenotype-associated protein families identified through attention analysis.

Overall, VenusPheno demonstrates that bacterial phenotype prediction can benefit from information distributed across multiple phenotype labels, particularly when labeled genomes are limited. More generally, this work considers a broader learning setting in which samples share a common input space and are associated with multiple, potentially interacting prediction targets. In such settings, relationships among labels can provide an additional source of supervision, but their value depends on whether the corresponding tasks reinforce or interfere with one another. Bacterial phenotype prediction provides a biologically important application in which to study this problem, while the underlying methodology is not restricted to this particular setting. The same principle could potentially be applied to other biological problems involving multiple labels and complex molecular inputs, such as predicting multiple clinical outcomes from sets of patient-derived cells or multiple ecosystem traits from microbial community profiles. By showing how task relationships can be learned from data and converted into reliable predictive gains, VenusPheno offers a general strategy for making more effective use of limited biological annotations.

## Notes

### Competing Interest Statement

The authors have declared no competing interest.

