## Supplementary Information for "VenusPheno: Transfer-Guided Task Grouping for Multi-Task Bacterial Phenotype Prediction"

### Supplementary Material

Jinling Yuan[1,2,3][0009−0006−1162−6967], Bingxin Zhou[2,3][0000−0002−3897−9766], and Liang Hong[2][0000−0003−0107−336X]

[1] School of Pharmaceutical Sciences, Shanghai Jiao Tong University, Shanghai 200240, China
[2] Institute of Natural Sciences, Shanghai Jiao Tong University, Shanghai 200240, China
[3] Equal contribution first authors.
{bingxin.zhou}@sjtu.edu.cn

#### A Supplementary Methods

This section describes dataset construction, genus-disjoint evaluation, benchmark protocols, VenusPheno training, and the analyses used for task transfer and biological interpretation.

##### A.1 Data sources, curation, and task definitions

Phenotype annotations and corresponding genome assembly accessions were obtained from the publicly available Bacformer dataset. Using these accessions, bacterial genome assemblies were retrieved from NCBI RefSeq and GenBank. Each genome assembly was treated as an individual sample and linked to its available phenotype annotations.

We constructed 14 binary classification tasks covering motility, sporulation, Gram staining, cell morphology, oxygen metabolism, and carbon source utilization. The original multiclass annotations for cell morphology and oxygen metabolism were converted into one-versus-rest binary tasks. Specifically, cell morphology was represented by bacillus and coccus, while oxygen metabolism was represented by aerobic, anaerobic, and facultative categories. For each binary task, samples belonging to the target category were assigned a positive label, while samples annotated with other categories of the same phenotype were assigned a negative label. The six carbon utilization tasks included citrate, glucose, glycerol, lactose, maltose, and sucrose. Only genomes with available annotations for a given task were included in its corresponding prediction dataset.

To obtain genus-level taxonomic assignments, we mapped the NCBI taxonomy identifiers (TaxIDs) associated with the genomes to their taxonomic lineages. Of the initial 12,338 genomes, 12,178 (98.7%) were successfully assigned to 2,061 distinct genera. The remaining 160 genomes without valid genus assignments were excluded from subsequent experiments.

##### A.2 Genus-disjoint data split

We adopted a genus-disjoint splitting strategy to evaluate model generalization to unseen bacterial genera. All genomes belonging to the same genus were assigned to the same partition, ensuring that no genus was shared across the training, validation, and test sets. The target split ratio was 70%, 10%, and 20%, respectively.

Because bacterial genera vary considerably in genome counts and phenotype distributions, random assignment at the genus level may lead to substantial differences in sample sizes and class distributions across partitions. To obtain a more balanced split while preserving genus-level separation, we performed a randomized search over 3,000 candidate partitions with a fixed random seed of 42.

For each genus, we calculated the total number of genomes and the numbers of positive and negative samples for each of the 14 tasks. Candidate partitions were evaluated according to their deviations from the target proportions in genome counts and task-specific label distributions. The optimization criterion combined the weighted mean absolute deviation with a penalty for the maximum deviation. Genome counts were assigned a weight of 8, compared with a weight of 1 for each task-label count, and the maximum-deviation penalty was weighted by 0.2. The candidate with the lowest score was selected as the final partition.

The resulting training, validation, and test sets contained 8,409, 1,257, and 2,512 genomes, corresponding to 1,443, 206, and 412 genera, respectively. We verified that the three partitions were genus-disjoint and examined task-specific positive and negative sample counts to ensure that each task could be evaluated in all partitions.

Supplementary Table S1 summarizes the definitions, sample sizes, class distributions, and genus counts of the 14 phenotype prediction tasks across the three partitions.

##### A.3 Benchmark methods and evaluation protocols

We compared VenusPheno with a range of existing bacterial phenotype prediction methods and machine learning approaches, including Traitar, PhenoTrex-XGB, BacML (RF, HGBM, RNN, and CNN), BacteriaNet, Bacformer, and Evo 2. These methods cover different strategies for representing bacterial genomes, including protein functional annotations, $k$-mer features, genomic sequences, and pretrained sequence embeddings.

All methods were evaluated using the same genus-disjoint training, validation, and test partitions. For each phenotype task, only genomes with available labels were included in the corresponding evaluation. Balanced accuracy (BAcc), calculated as the arithmetic mean of sensitivity and specificity, was used as the primary performance metric to account for differences in class distributions across tasks.

*Traitar.* Traitar was used as an annotation-based, non-retrained reference method. Protein-coding sequences predicted from the held-out test genomes were supplied to the official `traitar phenotype` workflow in `from_genes` mode. Traitar derives Pfam-based functional profiles from the predicted proteins and applies its published phenotype models to generate consensus phenotype calls. No labels from the genus-disjoint training or validation partitions were used to fit Traitar. Predictions were generated only for the union of held-out genomes and then matched to the task-specific test labels. We used the combined majority-vote phenotype predictions for evaluation.

*PhenoTrex-XGB.* For PhenoTrex, genome sequences from the fixed genus-disjoint partitions were converted into the PhenoTrex genotype representation using the official workflow. For each phenotype, the training and validation genomes were combined for the final model fit, whereas held-out test labels were not accessed during model training. We trained the package's default weighted XGBoost classifier separately for each task using `phenotrex train xgb` and generated test predictions using `phenotrex predict`. The reported binary calls and confidence scores were parsed from the native PhenoTrex output.

*BacML-RF.* The BacML random-forest baseline was trained independently for each phenotype using the fixed genus-disjoint partitions. Genomic DNA sequences were represented by $k$-mer count features using the BacML preprocessing workflow, and the random-forest classifier was fitted on the task-specific training genomes. Validation data were used for model-selection procedures and test genomes remained held out until final evaluation. Performance was summarized across three independent random-seed runs.

*BacML-HGBM.* The histogram gradient-boosting machine (HGBM) baseline used the same BacML $k$-mer count representation and the same task-specific genus-disjoint training, validation, and test partitions as BacML-RF. An HGBM classifier was trained separately for each phenotype. This baseline therefore evaluates a non-neural tree-ensemble model over genome-wide sequence composition features under the same split protocol. Performance was summarized across three independent random-seed runs.

*BacML-RNN.* For the BacML RNN baseline, each phenotype was modelled separately using the BacML recurrent neural-network architecture. Input genomes were encoded as one-hot DNA sequences using the BacML sequence-processing pipeline, with the corresponding $k$-mer configuration used in the benchmark implementation. Models were optimized with a learning rate of $3 \times 10^{-4}$ using the fixed training split; validation data were reserved for model selection, and the held-out genus-disjoint test partition was evaluated only after training. Performance was summarized across three independent random-seed runs.

*BacML-CNN.* The BacML CNN baseline was trained independently for each phenotype using one-hot encoded genomic DNA sequences and the BacML convolutional architecture. The same genus-disjoint train–validation–test assignments and preprocessing conventions as the other BacML baselines were retained. CNN models were trained with a learning rate of $3 \times 10^{-4}$, with validation data used for model selection. Performance was summarized across three independent random-seed runs.

*BacteriaNet.* BacteriaNet was evaluated as a pretrained genome-sequence phenotype predictor. For each held-out genome with an available nucleotide FASTA file, we applied the released `bacterianet predict` workflow using the provided pretrained model resources. Thus, BacteriaNet was not retrained on the VenusPheno labels or genus-disjoint training partition. Its predicted phenotypes were mapped to the corresponding benchmark tasks wherever a direct phenotype correspondence was available; tasks unsupported by the released predictor were not evaluated for this method.

*Bacformer.* Bacformer was evaluated using precomputed 480-dimensional genome-level embeddings. For each phenotype, we trained a classifier consisting of a multilayer perceptron with a 512-dimensional hidden layer, dropout of 0.3, and a binary output layer. Models were trained for up to 50 epochs using Adam (learning rate $10^{-4}$), with early stopping based on validation BAcc (patience 10). The best-performing model was evaluated on the held-out test genomes. Each experiment was repeated with three random seeds.

*Evo 2.* For Evo 2, we used precomputed 4,096-dimensional genome representations derived from pretrained Evo 2 sequence embeddings. For each phenotype, we trained an MLP classifier with a 512-dimensional hidden layer and dropout of 0.3. Models were trained for up to 80 epochs using AdamW (learning rate $10^{-4}$; weight decay $10^{-4}$), gradient clipping with a maximum norm of 1.0, and early stopping based on validation BAcc (patience 10). Each experiment was repeated with three random seeds.

##### A.4 VenusPheno training strategy

All experiments used predefined, mutually disjoint genus-level training, validation, and test partitions. The validation set was used exclusively for model selection and early stopping; the test set was evaluated only once using the selected checkpoint. For STL, the checkpoint with the highest validation BAcc was retained independently for each phenotype. For MTL, the retained checkpoint maximized macro-average validation BAcc across the 14-task panel. Binary predictions were generated using a fixed probability threshold of 0.5 for both validation and test evaluation.

For each binary task, positive-class weights were calculated from the training partition as the ratio of negative to positive samples and used in the binary cross-entropy loss. In MTL, phenotype labels absent for a genome were excluded using a task-specific label mask. Each observed task loss was first averaged over its valid samples before contributing to the joint objective. All reported results summarize three independently initialized runs.

Training used AdamW with a learning rate of $10^{-4}$ and weight decay of $10^{-4}$. Models were trained for at most 80 epochs with early stopping after 10 epochs without validation improvement. Gradients were clipped to an $\ell_2$ norm of 1.0. Automatic mixed-precision training was used on CUDA devices. No learning-rate scheduler was applied.

##### A.5 Protein annotation and pathway enrichment

For the attention-based analyses, high-attention proteins were annotated using eggNOG annotations, and pathway enrichment was assessed against KEGG pathways. Enrichment significance was adjusted for multiple testing using the Benjamini–Hochberg procedure; pathways displayed in the corresponding figure satisfy an adjusted FDR $\leq 0.05$. The protein-family attention comparison reports phenotype-positive minus phenotype-negative mean attention scores. Further analysis-specific sample counts and plotted quantities are provided in the figure captions.

#### B Supplementary Results

##### B.1 task transfer and grouping

**Complete task-transfer matrix** To further characterize task interactions under joint supervision, we examined the complete LOTO transfer matrix across all 14 phenotype prediction tasks (Supplementary Fig. S1). Among the 182 directed task pairs, 92 exhibited beneficial transfer and 89 exhibited detrimental transfer. The magnitude and direction of transfer varied substantially across source and target tasks, as reflected in their net outgoing and incoming transfer effects. Moreover, the transfer matrix revealed asymmetric relationships, indicating that the influence of one task on another was not necessarily reciprocal. These observations provide further evidence of heterogeneous task interactions under full-task MTL.

**Task grouping and sensitivity to $K$ and $\lambda$** To assess the sensitivity of task grouping to the penalty parameter, we compared partitions with three ($K = 3$) and four ($K = 4$) groups under $\lambda = 0.5$, 1, and 2 (Supplementary Fig. S2). For each fixed $K$, the same task assignments were obtained across all three penalty settings, indicating that the identified group structures were stable within the tested parameter range. At $\lambda = 1$, the three-group partition achieved a higher penalty-weighted transfer score than the four-group partition (43.71 versus 39.28 percentage points). As $\lambda$ increased, the objective scores decreased because detrimental transfer received a larger penalty, although the task assignments remained unchanged. At $\lambda = 2$, the four-group partition achieved a

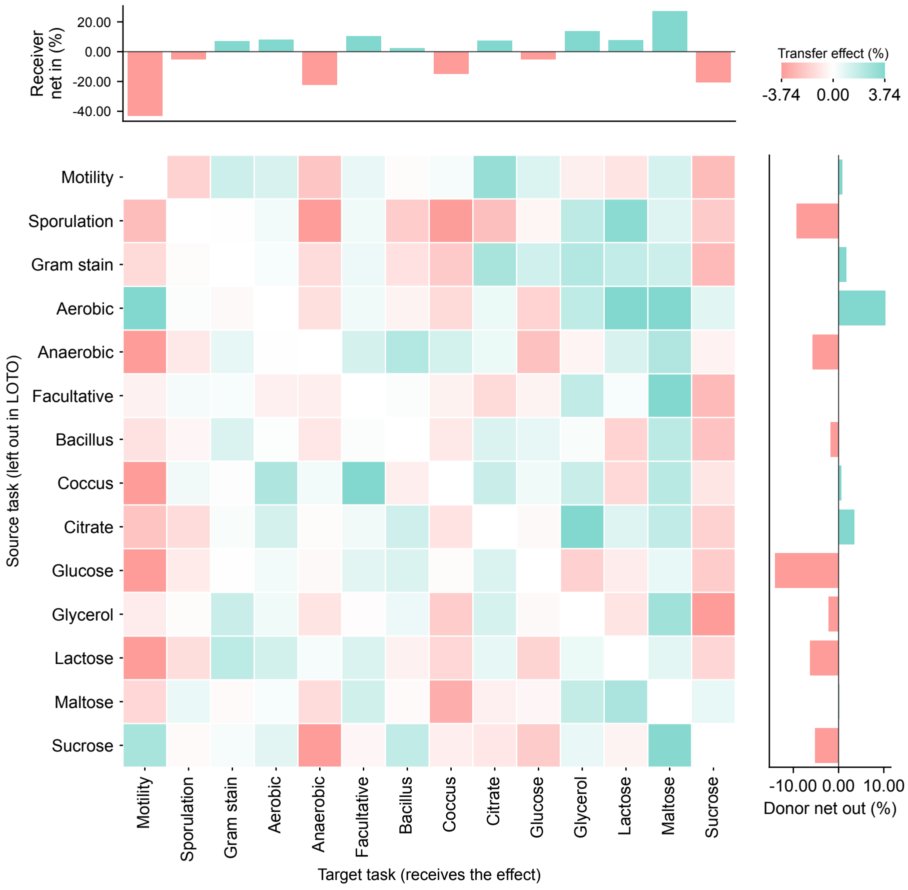


**Supplementary Figure S1. Directed task transfer relationships identified by leave-one-task-out (LOTO) analysis.** The heatmap shows pairwise transfer effects among 14 bacterial phenotype prediction tasks under full-task MTL. Rows represent source tasks excluded during LOTO, while columns represent target tasks receiving the transfer effect. Each entry corresponds to the difference in target-task BAcc between full-task training and training without the source task. Teal indicates beneficial transfer, whereas red indicates detrimental transfer. The top and right bar charts summarize the net incoming and outgoing transfer effects for each task, respectively. Diagonal entries are excluded. Transfer effects are expressed in percentage points.

slightly higher score than the three-group partition (32.82 versus 32.13 percentage points). Overall, these results indicate that the identified task memberships were insensitive to the tested penalty weights, while the relative scores of different group numbers depended on the penalty strength.

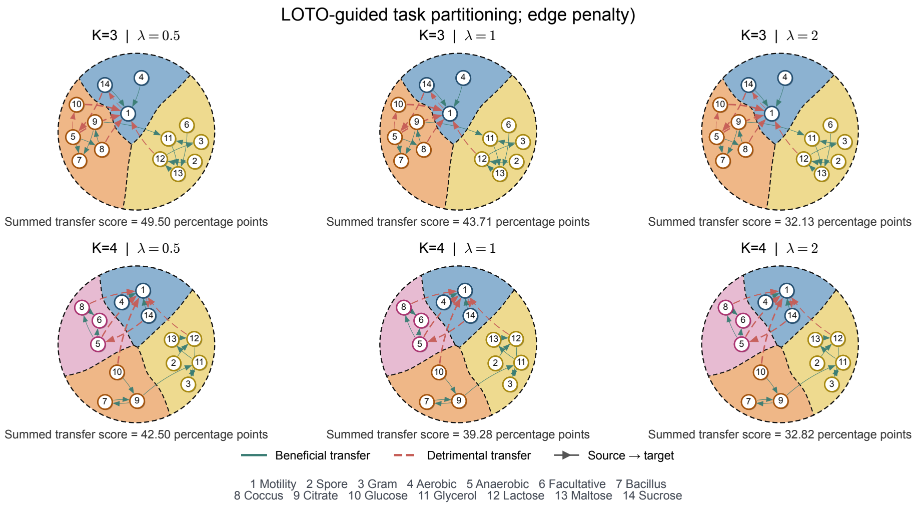


**Supplementary Figure S2. Sensitivity of LOTO-guided task partitioning to the negative-transfer penalty.** Directed task-transfer networks for $K = 3$ (top row) and $K = 4$ (bottom row) under three penalty settings ($\lambda = 0.5$, 1, and 2). Shaded regions indicate the identified task groups, and numbered nodes represent the 14 phenotype tasks as defined in the figure. Solid teal and dashed red arrows indicate beneficial and detrimental transfer, respectively, with arrow direction representing source-to-target transfer. The summed transfer score shown below each network corresponds to the penalty-weighted partitioning objective, expressed in percentage points. For each fixed $K$, group assignments remain unchanged across the tested penalty settings.

#### B.2 Biological interpretation

**High-attention proteins and pathway enrichment** This analysis was designed to examine protein-level model attention and functional enrichment in a held-out validation cohort; it was not intended to represent the full dataset. For each phenotype, we included all validation genomes with a positive task label for which task-matched STL attention weights had been successfully exported and the attention-weighted protein IDs could be strictly matched to the corresponding FAA protein records. All included genomes had eggNOG annotations available for functional analysis. This procedure yielded 31 motility-positive genomes and 25 sucrose-utilization-positive genomes, with all eligible positive validation genomes included. Thus, the cohort sizes reflect the number of positive genomes in the validation split for these tasks, rather than a manually selected subset.

Within each genome, proteins were ranked by their mean attention rank across the 16 PMA head-by-seed slots, and the top 10% were treated as high-attention proteins. For pathway enrichment, annotated high-attention proteins were compared with the other annotated proteins in the same positive genomes (Supplementary Fig. S3). For motility, FlgK was identified among the recurrent high-attention proteins in 29 of 31 genomes (94%). KEGG enrichment highlighted bacterial chemotaxis and flagellar assembly, with fold enrichments of approximately 6.5 and 6.1, respectively. RNA polymerase and the two-component system were also enriched.

For sucrose utilization, recurrent high-attention proteins included OppA (14/25 genomes, 56%), RbsR (12/25, 48%), and LacZ and MalL (11/25, 44% each). Enriched KEGG pathways included starch and sucrose metabolism and galactose metabolism, with fold enrichments of approximately 3.9 and 3.3, respectively. These results characterize attention patterns within the positive validation genomes and do not by themselves establish phenotype specificity.

**Protein-family attention under STL and grouped MTL** To further examine how joint supervision affects protein-level representations, we compared the distributions of family attention scores between phenotype-positive and phenotype-negative genomes under STL and grouped MTL (Supplementary Fig. S4). For motility, seven of the eight examined protein families exhibited larger positive-negative mean attention differences under grouped MTL. Particularly pronounced increases were observed for FliH, FliM, and flagellar hook proteins, whose differences increased from 0.10, 0.10, and 0.00 under STL to 0.55, 0.54, and 0.54 under grouped MTL, respectively. All eight motility-associated families showed significant positive-negative differences under grouped MTL after multiple-testing correction. For sucrose utilization, five of the six examined families exhibited increased attention differences, including PTS EIIA, PTS EIIB, PRD, and HTH_11 domains. However, these differences did not reach statistical significance after correction. Together, these distributions provide additional evidence that grouped MTL is associated with stronger phenotype-dependent attention patterns across several functionally relevant protein families, particularly those involved in bacterial motility.

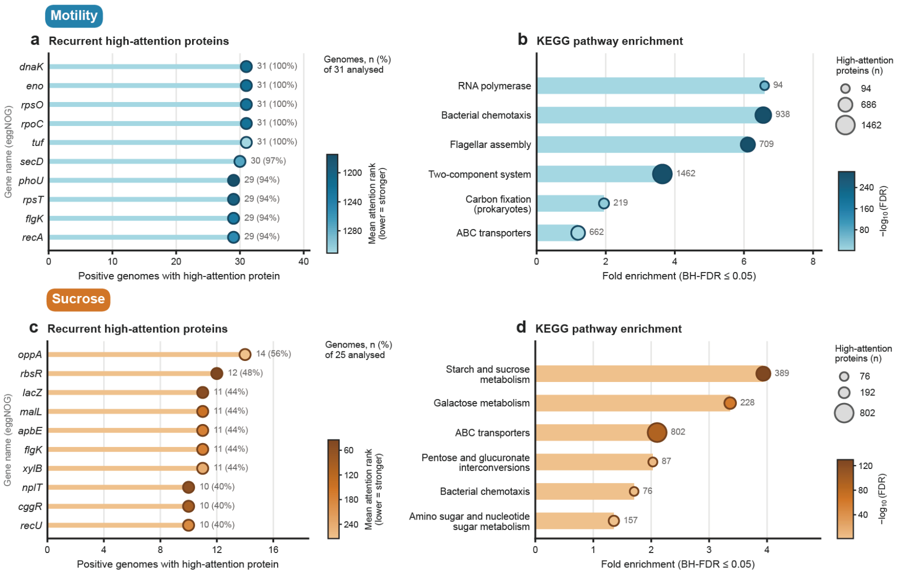


**Supplementary Figure S3. Recurrent high-attention proteins and KEGG pathway enrichment under STL.** (a, c) The ten most recurrent high-attention proteins identified in phenotype-positive genomes for motility (a) and sucrose utilization (c), based on eggNOG annotations. Horizontal bars indicate the number of positive genomes in which each protein annotation was identified among high-attention proteins, with counts and percentages shown on the right. Marker colors represent the mean attention rank, with lower ranks indicating stronger attention. The analyses include 31 motility-positive and 25 sucrose-positive genomes. (b, d) KEGG pathway enrichment of high-attention proteins for motility (b) and sucrose utilization (d). Horizontal bars represent fold enrichment, marker size indicates the number of high-attention proteins assigned to each pathway, and marker color represents enrichment significance ($-\log_{10}(\text{FDR})$). All displayed pathways satisfy the Benjamini-Hochberg adjusted significance threshold (FDR $\leq 0.05$).

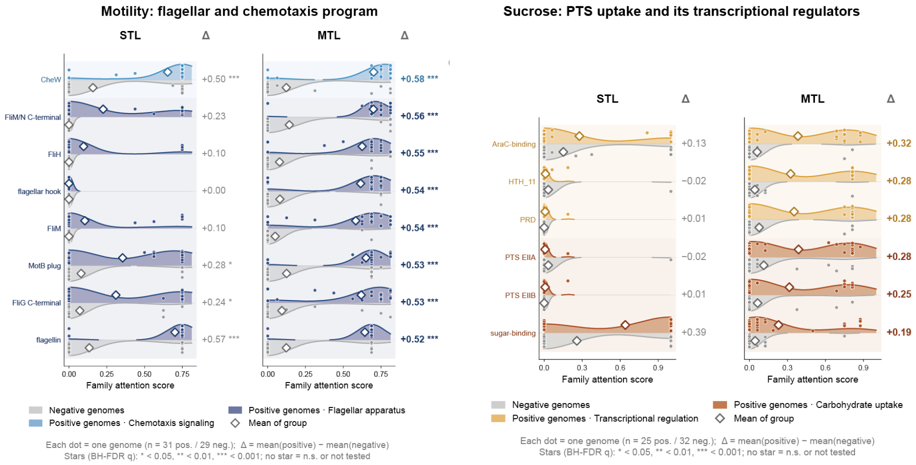


**Supplementary Figure S4. Protein-family attention distributions under STL and grouped MTL.** Attention-score distributions for eight motility-associated protein families (left) and six sucrose-utilization-associated protein families (right). For each family, gray and colored distributions represent phenotype-negative and phenotype-positive genomes, respectively. Colored distributions distinguish functional categories, including flagellar apparatus, chemotaxis signaling, carbohydrate uptake, and transcriptional regulation. Each dot represents an individual genome, and diamonds indicate group means. The attention contrast ($\Delta$) is calculated as the mean attention score of phenotype-positive genomes minus that of phenotype-negative genomes. Motility analysis includes 31 positive and 29 negative genomes, while sucrose-utilization analysis includes 25 positive and 32 negative genomes. Asterisks indicate statistical significance of positive-negative differences after Benjamini–Hochberg false discovery rate correction ($^{*}q < 0.05$, $^{**}q < 0.01$, $^{***}q < 0.001$).

**Supplementary Table S1.** Phenotype prediction tasks and genus-disjoint dataset splits. All 14 tasks are binary classification problems. For each split, the upper line gives the number of labeled genomes ($n$), and the lower gray line gives the numbers of positive and negative samples (#pos / #neg). Total $n$ is the number of labeled genomes per task, Pos. (%) is the overall proportion of positive samples, and Genera is the number of distinct genera represented in each task. The training, validation, and test splits contain disjoint sets of genera.

| Task | Train | Validation | Test | Total $n$ | Pos. (%) | Genera |
|---|---|---|---|---|---|---|
| *Cell structure* | | | | | | |
| Gram | 6,220 | 965 | 1,893 | 9,078 | 42.7 | 1,595 |
| | 2,548 / 3,672 | 348 / 617 | 978 / 915 | | | |
| Cell shape (bacillus) | 5,094 | 713 | 1,564 | 7,371 | 68.2 | 1,470 |
| | 3,419 / 1,675 | 496 / 217 | 1,115 / 449 | | | |
| Cell shape (coccus) | 5,094 | 713 | 1,564 | 7,371 | 20.0 | 1,470 |
| | 961 / 4,133 | 147 / 566 | 368 / 1,196 | | | |
| *Behavior & life cycle* | | | | | | |
| Motility | 6,341 | 985 | 1,855 | 9,181 | 42.5 | 1,634 |
| | 2,834 / 3,507 | 362 / 623 | 703 / 1,152 | | | |
| Sporulation | 5,866 | 798 | 1,748 | 8,412 | 16.1 | 1,578 |
| | 958 / 4,908 | 86 / 712 | 314 / 1,434 | | | |
| *Oxygen requirement* | | | | | | |
| Aerobic | 6,176 | 890 | 1,912 | 8,978 | 30.4 | 1,751 |
| | 1,910 / 4,266 | 225 / 665 | 598 / 1,314 | | | |
| Anaerobic | 6,176 | 890 | 1,912 | 8,978 | 14.8 | 1,751 |
| | 876 / 5,300 | 146 / 744 | 304 / 1,608 | | | |
| Facultative | 6,176 | 890 | 1,912 | 8,978 | 32.8 | 1,751 |
| | 1,953 / 4,223 | 339 / 551 | 654 / 1,258 | | | |
| *Carbon utilization* | | | | | | |
| Citrate | 2,204 | 286 | 538 | 3,028 | 15.6 | 1,156 |
| | 340 / 1,864 | 50 / 236 | 82 / 456 | | | |
| Glucose | 2,204 | 286 | 538 | 3,028 | 60.7 | 1,156 |
| | 1,361 / 843 | 151 / 135 | 325 / 213 | | | |
| Glycerol | 2,204 | 286 | 538 | 3,028 | 21.6 | 1,156 |
| | 469 / 1,735 | 70 / 216 | 115 / 423 | | | |
| Lactose | 2,204 | 286 | 538 | 3,028 | 25.6 | 1,156 |
| | 545 / 1,659 | 72 / 214 | 157 / 381 | | | |
| Maltose | 2,204 | 286 | 538 | 3,028 | 47.0 | 1,156 |
| | 1,056 / 1,148 | 123 / 163 | 244 / 294 | | | |
| Sucrose | 2,204 | 286 | 538 | 3,028 | 42.0 | 1,156 |
| | 917 / 1,287 | 124 / 162 | 232 / 306 | | | |

**Supplementary Table S2.** Architecture and training configurations for the matched STL and hard-sharing MTL models.

| **Shared configuration** | |
|---|---|
| Protein embeddings | ProtT5-XL-UniRef50 (1024 dimensions) |
| Genome encoder | ISAB followed by PMA |
| Attention configuration | 4 heads; 64 inducing points; 4 PMA seeds |
| Genome representation | Concatenated PMA outputs (4096 dimensions) |
| Activation / dropout | GELU / 0.3 |
| Orthogonality weight | $\lambda_{\text{orth}} = 0.01$ |
| Optimizer | AdamW (lr = $10^{-4}$, weight decay = $10^{-4}$) |
| Batch size | 32 |
| Gradient clipping | Global $\ell_2$ norm of 1.0 |
| Max. epochs / patience | 80 / 10 |
| Decision threshold | 0.5 |

| **Model-specific settings** | **STL** | **Hard-sharing MTL** |
|---|---|---|
| Prediction backbone | Independent residual MLP (512 hidden units) | Shared residual MLP (512 hidden units) |
| Classification heads | One binary linear head | 14 task-specific binary linear heads |
| Prediction loss | Class-weighted BCE | Masked, task-balanced weighted BCE |
| Checkpoint selection | Validation BAcc | Macro validation BAcc |
